# Sex differences in fear extinction are conditional, not categorical

**DOI:** 10.64898/2026.10.27.762318

**Authors:** Minh Nguyen, Angelos Didachos, Benno Roozendaal, Marloes J.A.G. Henckens, Kubra Gulmez Karaca

## Abstract

Threat memories are dynamic, and their behavioral expression can change with subsequent experience and the passage of time. Extinction training can reduce conditioned fear responses, yet these responses may re-emerge over time through spontaneous recovery. Although biological sex has been implicated as modulatory factor in fear and extinction memory processes, studies have reported inconsistent findings. Here, we investigated how differences in extinction training and the timing of memory assessment influence extinction memory in male and female mice. Using contextual fear conditioning, we assessed fear acquisition and threat retention, characterized within-session and across-session extinction during repeated training, and compared shorter (3-minute) and longer (6-minute) daily extinction sessions. We subsequently evaluated extinction memory retention and spontaneous recovery at a later time point. Estrous-cycle stage in females was monitored daily throughout the experimental protocol to examine its potential association with fear and extinction behavior. We found that males and females demonstrated comparable fear acquisition and threat memory, as well as similar extinction progression across days. During longer extinction sessions, but not shorter, males showed greater within-session reductions in freezing, whereas females exhibited modestly higher overall freezing. We found no significant sex difference in the magnitude of spontaneous recovery. There was no significant association between estrous-cycle stage and freezing behavior. Together, our findings suggest that sex-dependent differences in contextual fear extinction are not uniformly expressed, but rather depend on the experimental conditions under which extinction is assessed.

**Highlights:**

- Male and female mice showed similar threat memory acquisition and expression.
- Extinction progression across days was comparable in both sexes.
- Males showed greater within-session freezing reductions in longer extinction trials.
- Spontaneous fear recovery did not differ significantly between sexes.
- Freezing behavior did not depend on estrous-cycle stage.

## 1. Introduction

Threat memories are not static representations of past experiences but remain dynamic. The behavioral expression of fear can be influenced by several factors, including the circumstances of fear acquisition and retrieval, subsequent experiences of the individual, reminders of the original event, and the passage of time itself (Maren et al., 2013). Fear extinction provides a well-established example of how new experiences can alter the expression of previously acquired threat memories over time. During extinction learning, repeated exposure to a conditioned stimulus without reinforcement typically leads to a progressive reduction in conditioned fear responses (Quirk and Mueller, 2008). However, this reduction is not necessarily permanent, as extinguished fear responses can re-emerge with the passage of time, a phenomenon known as spontaneous recovery of fear (Maren et al., 2013; Rescorla, 2004). Thus, even after extinction, the expression of conditioned fear may be subject to change over time.

Beyond experience and time, biological sex has been extensively investigated as a potential factor influencing the formation and expression of threat and extinction memories. This interest is partly driven by the higher prevalence of certain stress- and fear-related disorders, including post-traumatic stress disorder (PTSD) and generalized anxiety disorder, among women compared with men (Kornfield et al., 2018; Olff, 2017; Riccardi et al., 2024). However, findings on sex differences in fear acquisition and extinction remain inconsistent (Bauer, 2023; Velasco et al., 2023). Some studies have reported greater fear expression or impaired extinction in females (Baran et al., 2009; Matsuda et al., 2015), whereas others found comparable performance between sexes (Schoenberg et al., 2022; Voulo and Parsons, 2017) or even faster extinction in females (Riccardi et al., 2024). A recent meta-analysis found no overall sex differences in freezing behavior, the main behavioral proxy of fear in rodents, before or after extinction, but a substantial heterogeneity across studies and a strong male bias in rodent literature (Sun et al., 2026). Fluctuations in ovarian hormones across the estrous cycle have been proposed as one possible source of variability in fear and extinction learning and memory (Hohorst et al., 2026; Milad et al., 2009; Peyrot et al., 2024), although the extent to which estrous-cycle stage accounts for differences in behavioral outcomes remains unclear. Furthermore, sex differences in the longer-term persistence of extinction memory and spontaneous recovery have received relatively little attention (Matsuda et al., 2015; Park et al., 2017). Together, these findings raise the possibility that sex-dependent effects vary with the conditions under which fear memories are acquired, extinguished, and retrieved. Yet, these factors are often investigated separately, limiting our understanding of how they shape threat memory expression across different experiences and time intervals.

Here, we investigated how three major sources of variability, i.e., biological sex, extinction experience, and the timing of memory assessment, relate to the expression and persistence of threat and extinction memories. Using contextual fear conditioning in male and female mice, we assessed fear acquisition and threat memory recall, characterized within-session and across-session extinction during repeated training, and compared different daily extinction session durations (either 6-min or 3-min). We subsequently evaluated extinction memory retention (3-day) and spontaneous recovery at a later time point (2-3 week). In female mice, we also monitored estrous-cycle stage daily, from initial handling through the final memory test, to examine its potential association with fear expression. By investigating these factors within a common experimental framework, we aimed to determine how extinction experience and memory age relate to fear expression and persistence of extinction memory, and whether sex-dependent differences are consistently observed across these conditions.

## 2. Materials and Methods

### 2.1. Animals and housing

Male and female C57BL/6J mice were obtained from Charles River Laboratories (France). Mice were 6-8 weeks old at the beginning of the experiment and were housed in pairs in standard cages at the Central Animal Facility of Radboudumc (Nijmegen, The Netherlands). Housing conditions were maintained on a 12-h light/dark cycle (lights on at 07:00 h; lights off at 19:00 h), at 22°C, approximately 72% relative humidity, and a light intensity of 47 lux. Food and water were available ad libitum. Behavioral experiments were conducted between 08:00 and 13:00 h.

A total of 44 mice (22 males, 22 females) were initially included in the study. One animal reached a humane endpoint, and one male was removed from the study because he jumped onto the chamber lid and missed two footshocks. One male was excluded from the contextual fear conditioning analysis because the video was not available due to a technical problem. Final group sizes were 42 mice for extinction and tests (20 males, 22 females) and 41 for CFC (19 males, 22 females). All experimental procedures were performed in accordance with Dutch legislation concerning animal welfare and relevant international guidelines and were approved by the Central Committee for Animal Experiments (The Hague, The Netherlands; approval number 2022-0020).

### 2.2. Contextual fear conditioning

During habituation phase, mice were gently handled for 3-5 min daily over four consecutive days in the experimental room. Animals were then left undisturbed for one day before conditioning. Contextual fear conditioning (CFC) was performed in a chamber (Campden Instruments, UK) equipped with white, rounded walls, a stainless-steel grid floor, and a metal tray underneath. Each mouse received five footshocks (1 mA, 2 s each) delivered at 178, 208, 238, 268, and 298 s after the beginning of the session. Thirty seconds after the final footshock, mice were returned to their home cages. The chamber was cleaned with 70% ethanol between animals. No auditory cues were presented during conditioning or subsequent behavioral testing. All sessions were video-recorded for behavioral analysis.

### 2.3. Extinction training and memory testing

Extinction training began three days after CFC and was conducted over five consecutive days. During each extinction session, mice were returned to the conditioning context without footshock delivery. Animals were assigned to either a short-duration (3 min/day) or long-duration (6 min/day) extinction protocol. Following each session, mice were returned to their home cages, and the conditioning chamber was cleaned with 70% ethanol between animals. Extinction memory retention was assessed three days after the final extinction session. Mice were re-exposed to the conditioning context without footshocks for either 3 or 6 min, matching their extinction-training protocol. Spontaneous recovery of conditioned fear was subsequently assessed by re-exposing mice to the conditioning context without footshocks. Spontaneous recovery was assessed 2-3 weeks after the final extinction session. No footshocks or auditory cues were presented during extinction training or memory testing.

### 2.4. Automated behavioral quantification

Behavior was recorded using overhead cameras positioned above the conditioning chambers at 25 frames per second. Freezing was defined as the absence of movement except for movements necessary for respiration. Freezing was quantified using DeepLabCut (DLC; (Mathis et al., 2018)) and EasyFreezy, a Python-based behavioral analysis tool developed by a co-author of this study (https://github.com/AngelosDid/EasyFreezy).

A DeepLabCut network was trained to track the nose, left ear, right ear, and midpoint between the ears. The training dataset comprised 12 videos, each 3 or 6 min long, recorded at 960 × 576 pixels. Twenty frames per video were manually labeled, with 95% allocated to training. A ResNet-50 network was trained through three rounds of iterative training and refinement, yielding a test error of 3.31 pixels. Freezing was detected using EasyFreezy based on displacement of the midpoint between the ears, with an additional nose-movement filter to reduce false-positive detections. A minimum duration of 12 consecutive frames (0.48 s) was required to classify a freezing episode. Freezing was expressed as the percentage of time spent freezing within each analysis interval. To evaluate the performance of automated freezing detection, eight videos from two experimental batches (four male and four female mice) were manually scored using Behavioral Observation Research Interactive Software (Friard and Gamba, 2016). Freezing percentages obtained from manual and automated scoring were strongly correlated (Pearson’s *r* = 0.963, n = 8, *p* < 0.001). Automated freezing measurements were used for subsequent behavioral analyses.

Within-session freezing was analyzed minute by minute and by comparing the beginning and end of each extinction session. For the 6-minute protocol, freezing during the first and last 3 min was compared. For the 3-minute protocol, freezing during the first and last minute was compared.

### 2.5. Estrous-cycle monitoring

Estrous-cycle stage was assessed using vaginal cytology in female mice. Samples were collected with demineralized-water-moistened swabs applied to the vaginal opening. Male mice underwent a corresponding swabbing procedure at the anal opening to provide comparable handling-related stimulation. During acclimatization (1 week prior to the habituation) and handling, samples were collected between 09:00 and 12:00 h. On conditioning, extinction, and memory-testing days, sampling was performed approximately six hours after behavioral procedures, between 15:00 and 18:00 h, not to interfere with the early consolidation window. Collected samples were transferred onto a drop of demineralized water on Superfrost microscope slides (catalog no. 125447, Thermo Fisher Scientific, The Netherlands), air-dried, and stained with Giemsa solution for 10-12 min at room temperature. The staining solution was prepared using Giemsa’s azur eosin methylene blue solution (catalog no. 109204, Sigma, Germany) diluted in Tris-HCl buffer (pH 7.0-7.4). Following staining, slides were allowed to dry. Images were acquired using a Leica DM2000 bright-field microscope (Leica Microsystems, Germany) equipped with a digital camera and a 10× objective. Estrous-cycle stages were classified manually based on the relative abundance and morphology of leukocytes, nucleated epithelial cells, and cornified epithelial cells, following established criteria (Ajayi and Akhigbe, 2020; Byers et al., 2012; Cora et al., 2015). Stages were categorized as proestrus (P), estrus (E), metestrus (M), or diestrus (D). Estrous stages were grouped into P/E and M/D to facilitate comparisons between broader phases of the estrous cycle while maintaining sufficient sample sizes for analysis.

### 2.6. Statistical analysis

Statistical analyses were performed using GraphPad Prism for Windows (Version 11.0.2). Data are presented as mean ± SEM, and statistical significance was set at *p* < 0.05. Contextual fear acquisition, across-session extinction, and extinction memory retention and spontaneous recovery were analyzed using two-way repeated-measures ANOVA or mixed-effects models, as appropriate. Within-session freezing dynamics were analyzed using three-way repeated-measures ANOVA or mixed-effects models, while within-session freezing reductions were analyzed using two-way repeated-measures ANOVA.

Comparisons between extinction training durations were performed using three-way mixed-effects analysis. Unpaired two-tailed *t*-tests, with Welch’s correction where appropriate, were used for comparisons between sexes, and one-sample *t*-tests were used to assess changes relative to zero. Freezing was compared between P/E and M/D females at each session using unpaired Welch’s *t*-tests, corrected for multiple comparisons across sessions (Benjamini, Krieger and Yekutieli; Q = 1%). Multiple comparisons were performed using Sidak’s, Tukey’s, or Fisher’s LSD tests, as specified in the figure legends.

## 3. Results

### 3.1. Male and female mice exhibited comparable contextual fear acquisition

We first examined whether male and female mice differed in their acquisition of contextual fear (Fig. 1A). During conditioning, both sexes initially displayed low levels of freezing, followed by a progressive increase upon repeated footshock delivery (Fig. 1B). A two-way repeated measures ANOVA revealed a significant effect of time (*F*(3.00, 117.0) = 193.8, *p* < 0.0001), but no significant effect of sex (*F*(1, 39) = 2.464, *p* = 0.1246) or time × sex interaction (*F*(3.00, 117.0) = 1.976, *p* = 0.1213; two-way repeated-measures ANOVA). Together, these findings indicate that male and female mice acquired contextual fear with no detectable sex differences in freezing during conditioning.

**Figure 1.**
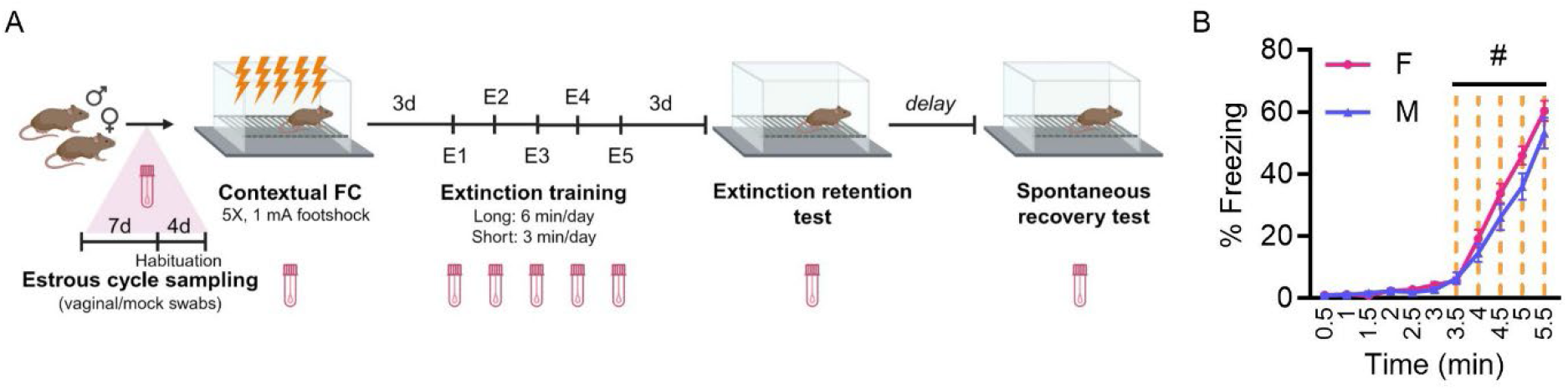
Experimental design and contextual fear acquisition in male and female mice. **A)** Schematic representation of the experimental protocol. During habituation phase, mice were gently handled for 3-5 min daily over four consecutive days in the experimental room. Male and female mice underwent contextual fear conditioning, followed three days later by five consecutive days of extinction training (E1-E5) consisting of daily re-exposure to the conditioning context without footshock, either for 6 min or 3 min per day. Extinction retention (3-day after final extinction session) and spontaneous fear recovery (2-3 week after final extinction session) were subsequently assessed. **B)** Percentage freezing during contextual fear conditioning, including the baseline period and successive footshock presentations. Male: *n* = 19; female: *n* = 22. Data are presented as mean ± SEM. Figure was generated using BioRender.

### 3.2. Sex differences in within-session, but not across-session, fear extinction depended on extinction training duration

Next, we investigated whether male and female mice differed in fear extinction and whether these differences varied with extinction training conditions. To this end, three days after fear conditioning, mice were re-exposed to the conditioning context for 6 minutes daily over five consecutive days, without footshock exposure. We first examined freezing during the initial extinction session (E1), which also served as an assessment of threat memory recall three days after conditioning. Freezing levels were comparable between males and females (38.64% and 42.43%, respectively; unpaired Welch’s *t*-test, *t*(22.80) = 0.8633, *p* = 0.3970), indicating no detectable sex differences in threat memory expression at this 3-day retention test. Across subsequent extinction sessions, freezing progressively decreased in both sexes, with significantly lower levels on each extinction day compared with the first (E1 vs. E2, *p* = 0.0005; E1 vs. E3-E5, all *p* < 0.0001; Sidak’s multiple comparisons test; Fig. 2A). A mixed-effects analysis revealed a significant effect of session (*F*(2.553, 63.19) = 29.31, *p* < 0.0001) and a modest, yet significant, effect of sex (*F*(1, 25) = 4.269, *p* = 0.0493), but no session × sex interaction (*F*(2.553, 63.19) = 0.3938, *p* = 0.7259). These findings indicate that despite higher overall freezing levels in females across extinction sessions, both sexes exhibited a comparable fear extinction across days.

**Figure 2.**
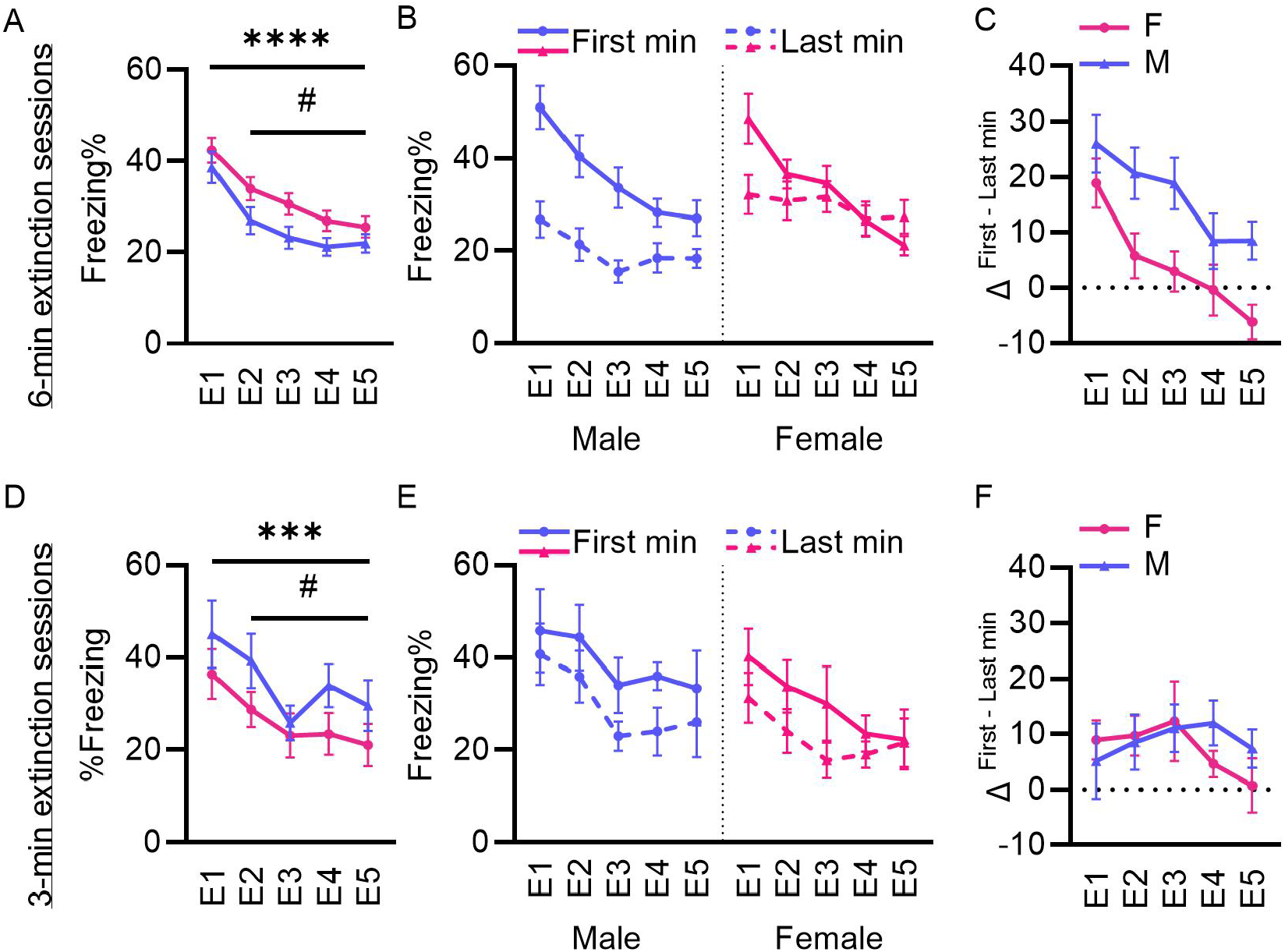
Within- and across-session fear extinction in male and female mice under different training durations. **A)** Percentage of f reezing across f ive consecutive days of 6-minute extinction training (E1-E5), beginning three days af ter contextual f ear conditioning. Freezing decreased signif icantly across sessions in both sexes, with higher overall f reezing levels in f emales (n = 14) than males (n =13) **B)** Percentage of f reezing during the f irst and last minutes of each 6-minute extinction session. **C)** Within-session f reezing reduction, calculated as the delta (Δ) in percentage of f reezing during the f irst minute minus last minute of each 6-minute extinction session. **D)** Percentage of f reezing across f ive consecutive days of 3-minute extinction training in a separate cohort of f emale (n = 8) and male (n = 7) mice. Freezing decreased signif icantly across sessions, without detectable sex diff erences. **E)** Percentage f reezing during the f irst and last minute of each 3-minute extinction session. **F)** Within-session f reezing reduction, calculated as the delta (Δ) in percentage of f reezing during the f irst minute minus that during the last minute of each 3-minute extinction session. Data are presented as mean ± SEM.

We next examined within-session freezing dynamics by comparing the first and last minute of each extinction session to determine whether males and females differed in the immediate reduction of fear expression in each training session, beyond the changes observed across consecutive days of extinction training (Fig. 2B). A three-way mixed-effects analysis revealed a significant effect of time (*F*(1, 26) = 31.13, *p* < 0.0001) and session × time (*F*(4, 104) = 7.137, *p* < 0.0001), indicating that freezing levels differed between the beginning and end of extinction session and that the magnitude of these changes varied across extinction days. Importantly, a significant time × sex interaction was observed (*F*(1, 26) = 12.17, *p* = 0.0017), indicating that within-session freezing changes differed between sexes, consistently across extinction days (session × time × sex: *F*(4.104) = 0.3249, *p* = 0.8607). To further characterize these sex-differences in within-session freezing levels, we calculated the reduction in freezing between the first and last minute of each session (Fig. 2C). Males exhibited greater within-session freezing reductions than females (main effect of sex: *F*(1, 25) = 12.03, *p* = 0.0019; two-way repeated-measures ANOVA). Together, these results indicate that, despite a similar progression of extinction across days, males showed greater within-session freezing reductions than females, and more robust threat memory expression in females during 6-minute extinction training.

We then investigated whether these sex differences were conserved when the duration of daily extinction training was shortened. A separate group of mice underwent five consecutive days of 3-minute exposure to the conditioning context without footshock, reducing the duration of each exposure by half. As in the 6-minute protocol, freezing progressively decreased across extinction days in both sexes (Fig. 2D), reflected by a significant main effect of session (*p* = 0.0006; two-way repeated-measures ANOVA). Freezing was significantly lower than on the first extinction day at all subsequent sessions (E1 vs. E2, *p* = 0.0496; E1 vs. E3, *p* < 0.0001; E1 vs. E4, *p* = 0.0214; E1 vs. E5, *p* = 0.0100), indicating that the shorter training protocol produced effective extinction across repeated sessions. Notably, with this protocol, the main effect of sex as observed for the extended 6-minute extinction protocol, was not replicated (*p* = 0.1867). There was also no significant session × sex interaction (*p* = 0.6605).

To assess within-session extinction during the shorter protocol, we compared freezing during the first and last minute of each session (Fig. 2E). A three-way repeated-measures ANOVA revealed significant effects of time (*p* = 0.0004) and session (*p* = 0.0022), indicating changes in freezing within and across extinction sessions. In contrast to the 6-minute protocol, neither the time × sex interaction (*p* = 0.6590) nor the session × time × sex interaction (*p* = 0.6013) was significant. Consistent with these findings, analysis of the reduction in freezing between the first and last minute revealed no significant main effect of sex (*F*(1, 13) = 0.2039, *p* = 0.6590), session (*F*(2.969, 38.59) = 0.7587, *p* = 0.5229), or session × sex interaction (*F*(2.969, 38.59) = 0.6272, *p* = 0.6002; Fig. 2F). Thus, although freezing decreased within sessions, the shorter extinction protocol did not reveal detectable sex differences in these within-session extinction levels.

Altogether, these findings indicate that sex differences in contextual fear extinction were more apparent in within-session freezing dynamics than in the progression of extinction across days. Greater within-session freezing reductions were observed in males during 6-minute extinction training, whereas no detectable sex differences emerged during 3-minute sessions. These findings suggest that the detection of sex-dependent differences in within-session freezing may vary with extinction training duration.

### 3.3. Sex differences in within-test freezing dynamics do not translate into differences in spontaneous fear recovery

We next investigated whether males and females differed in the retention of extinction memory and its persistence over time. To this end, we compared freezing during the extinction memory test (ExT), conducted three days after the final extinction training session (E5) of the 6-min extinction protocol, with freezing during the subsequent spontaneous recovery test (SR), conducted several weeks later (Fig. 3A). A two-way repeated-measures ANOVA revealed a significant effect of test (*p* = 0.0035), indicating an overall increase in freezing between ExT and SR, consistent with spontaneous fear recovery over time. Post hoc comparisons revealed a significant increase in freezing in females (*p* = 0.0049), but not in males (*p* = 0.1440). However, there was no significant effect of sex (*p* = 0.5587) or test × sex interaction (*p* = 0.3016), suggesting no difference in the magnitude of spontaneous recovery between sexes.

**Figure 3.**
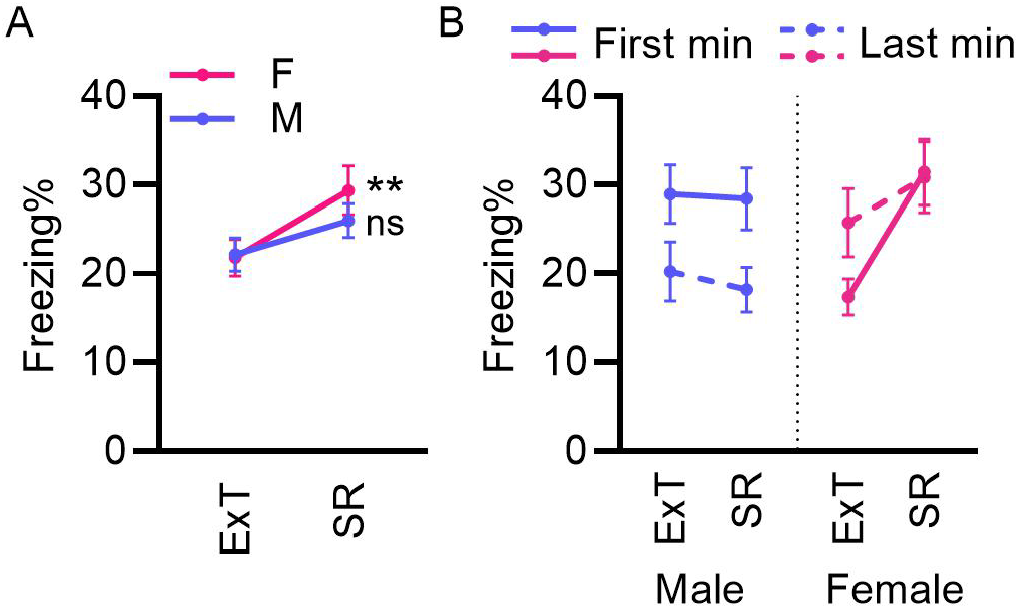
Extinction memory retention and spontaneous fear recovery in male and female mice. **A)** Percentage of freezing in female (n= 14) and male (n = 13) mice during the extinction memory test (ExT), conducted three days after the final extinction training session, and the subsequent spontaneous recovery test (SR), conducted several weeks later. **B)** Percentage freezing during the first and last minute of ExT and SR. Data are presented as mean ± SEM in (A) and (C), and as individual values with mean ± SEM in (B). ExT, extinction memory test; SR, spontaneous recovery test.

To assess whether sex-related differences in within-session freezing dynamics were also present during the ExT and SR tests, we compared freezing during the first and last minute of each test (Fig. 3B). A three-way repeated-measures ANOVA revealed significant test × sex (*p* = 0.0385) and time × sex (*p* = 0.0308) interactions, indicating that differences in freezing across tests and within-test time points varied between males and females. When considering the first and last minutes, the main effects of test (*p* = 0.0541), time (*p* = 0.3972), and sex (*p* = 0.3121), as well as the test × time (*p* = 0.0742) and test × time × sex (*p* = 0.4030) interactions, were not significant. These findings suggest that males and females differ in the temporal dynamics of freezing during memory testing, but there was no significant evidence that sex-dependent within-session changes differed between ExT and SR or translated into detectable sex differences in spontaneous fear recovery assessed across the full testing sessions.

Finally, we examined whether estrous stage was associated with freezing behavior across the different phases of the experiment. For this, estrous cycle stage was monitored throughout the experiment, and females were classified as being in proestrus/estrus (P/E) or metestrus/diestrus (M/D) at each behavioral assessment, following previous fear-extinction studies that grouped these stages according to their broadly higher versus lower ovarian hormone profiles (Bouchet et al., 2017; Hohorst et al., 2026). Freezing did not differ significantly between P/E and M/D females during contextual fear conditioning, extinction training (E1-E5), extinction memory testing or spontaneous recovery, nor did the within-session reduction in freezing in either extinction cohort (Supplementary Fig. 1; Welch’s *t*-tests per session, false discovery rate-corrected across sessions within each panel, all *q* ≥ 0.15). It should be noted, however, that due to the small sample sizes in the M/D groups, the analyses were primarily sensitive to large estrous-stage-related effects.

## 4. Discussion

Our findings indicate that sex differences in fear expression and extinction memory are not uniformly observed, but may depend on the conditions under which extinction training is conducted, the memory test examined, and whether fear behavior is characterized by initial fear expression, overall freezing levels, or within-session changes in freezing. Male and female mice showed comparable acquisition of contextual fear and a similar overall progression of extinction across repeated training sessions. However, during longer extinction sessions, males showed greater within-session reductions in freezing, whereas females exhibited higher overall freezing levels. These differences were not detected during shorter extinction sessions. Furthermore, although males and females differed in within-test freezing dynamics, no significant sex differences were observed in the magnitude of spontaneous fear recovery. Together, these findings suggest that sex differences in fear and extinction memories are conditional rather than categorical, and that differences in immediate fear expression do not necessarily translate into differences in the persistence of extinction memory. By examining biological sex, extinction experience, and memory assessment intervals within a common experimental framework, our study highlights the importance of distinguishing behavioral expression of fear during extinction training from memory persistence when interpreting sex-dependent differences in fear extinction.

Previous studies investigating sex differences in fear extinction have reported impaired extinction in females (Baran et al., 2009; Voulo and Parsons, 2017), faster extinction in females (Riccardi et al., 2024), or differences depending on hormonal state (Hohorst et al., 2026; Milad et al., 2009; Peyrot et al., 2024) or the interval between fear conditioning and extinction training (Schoenberg et al., 2022), while others found no sex differences ((Schoenberg et al., 2022) for recent memories). These inconsistencies are further complicated by a strong male bias in preclinical research. A recent meta-analysis reported that more than 70% of mouse and 87% of rat extinction studies included only males, and that those studies comparing sexes found no overall differences in freezing before or after extinction, despite substantial heterogeneity (Sun et al., 2026). Our findings suggest that some of this variability may be caused by potentially different aspects of extinction behavior being examined. Although males and females showed comparable progression of extinction across days, females exhibited modestly higher overall freezing during longer extinction training, whereas males showed greater within-session freezing reductions. Thus, even when freezing is used as the common behavioral measure, analyses of overall freezing levels, across-session extinction, and within-session changes may lead to different conclusions regarding sex differences.

Consistent with this, sex-dependent differences in within-session freezing dynamics have been reported during both 20-min long cued (Marin-Blasco et al., 2024) and 20-min long contextual fear extinction sessions (Matsuda et al., 2015). In our study, sex differences in within-session freezing reductions were detected during longer, but not shorter, extinction sessions, suggesting that training conditions and measurement windows may influence the detection of sex-dependent effects. Beyond extinction training, the timing of memory assessment may also be important, as extinguished fear can re-emerge with the passage of time (Quirk and Mueller, 2008; Rescorla, 2004). While Marin-Blasco et al. (2024) did not assess subsequent extinction memory retention or spontaneous recovery (Marin-Blasco et al., 2024), Matsuda et al. (2015) found that females required more extinction sessions than males to prevent spontaneous recovery (Matsuda et al., 2015). Here, we found that freezing increased significantly between extinction memory testing and spontaneous recovery in females but not males. However, neither the test × sex interaction nor the direct comparison of recovery magnitude revealed a significant sex difference, indicating that the increase observed in females alone does not establish greater spontaneous recovery in females. Thus, sex differences in immediate freezing reductions were not accompanied by detectable differences in subsequent spontaneous recovery. By examining extinction training, memory retention, and spontaneous recovery within the same experimental framework, our study extends previous work by distinguishing between immediate fear expression and longer-term extinction memory.

These findings may also have implications for extinction-based interventions, such as exposure therapy for anxiety- and trauma-related disorders. Although reductions in fear during extinction training are often considered an indication of successful extinction, they may not reflect the strength or persistence of extinction memory (Craske et al., 2008; Craske and Mystkowski, 2007). In our study, males showed greater within-session reductions in freezing during longer extinction sessions. These differences were not accompanied by a detectable sex difference in subsequent spontaneous recovery. Similar distinctions have been reported in clinical studies. In patients with PTSD, 30-minute imaginal exposure sessions produced less within-session habituation than 60-minute sessions, without a significant difference in treatment outcomes (Van Minnen and Foa, 2006). Moreover, a randomized trial found that 60-minute prolonged exposure sessions were not inferior to 90-minute sessions (Foa et al., 2022). Together, these findings suggest that session duration may influence immediate fear reduction without necessarily determining longer-term outcomes. Although contextual fear extinction in mice cannot directly predict clinical treatment responses, our findings highlight the importance of assessing longer-term fear recovery rather than relying solely on immediate fear reduction, while also considering how the duration of extinction training may affect the detection of sex differences.

Ovarian hormones have been implicated in the regulation of fear extinction. In rats, females in proestrus/estrus showed lower freezing during extinction than males and females in metestrus/diestrus (Hohorst et al., 2026), and extinction learned during proestrus was better consolidated than extinction learned during metestrus (Milad et al., 2010, 2009). Similarly, women tested during the mid-cycle phase showed better extinction retention than those tested during the early follicular phase (Peyrot et al., 2024). One strength of our study is that estrous stage was monitored throughout the experimental protocol, from fear conditioning through repeated extinction sessions and subsequent memory assessments, allowing us to examine potential stage-related differences at multiple stages of fear and extinction memory processing. However, we did not detect significant differences in freezing between females in proestrus/estrus and metestrus/diestrus. Admittedly, the small sample sizes in the metestrus/diestrus groups meant that our analyses were primarily sensitive to large estrous-stage-related effects. Moreover, we did not directly measure circulating hormone concentrations. Thus, although our findings did not reveal a significant association between estrous stage and fear extinction, more subtle hormonal influences cannot be excluded.

In conclusion, our findings show that sex differences in contextual fear extinction are not uniform, but depend the aspect of fear expression examined (initial freezing, whole-session freezing, or within-session changes) and the conditions under which extinction is assessed. Differences in fear expression during extinction training were not accompanied by detectable sex differences in the longer-term persistence of extinction memory. Whether these behavioral patterns reflect sex-dependent differences in the neural processing of fear and extinction memories, particularly within the neuronal ensembles that encode and represent these memories (Gulmez Karaca et al., 2021; Josselyn et al., 2015; Kupke and Oliveira, 2025), remains to be investigated. Understanding how biological sex, extinction experience, and memory assessment timing relate to these underlying neural representations may provide new insights into the mechanisms governing fear extinction and memory persistence.

## Acknowledgements

We thank Pleuni Schreurs for technical assistance throughout the project and the Radboudumc Technology Center Animal Research Facility for technical support with animal experiments. Schematic illustrations were created using BioRender.com.

## Author contributions

KGK, MJAGH, and BR conceptualized the study, discussed the data, and revised the manuscript. KGK led the study, designed and performed the experiments, analyzed the data, supervised the project, and wrote the original manuscript draft. MN performed experiments and analyzed data. AD developed the EasyFreezy software used for automated freezing analysis and provided training on its use.

## Supplementary Figures

**Supplementary Figure 1.**
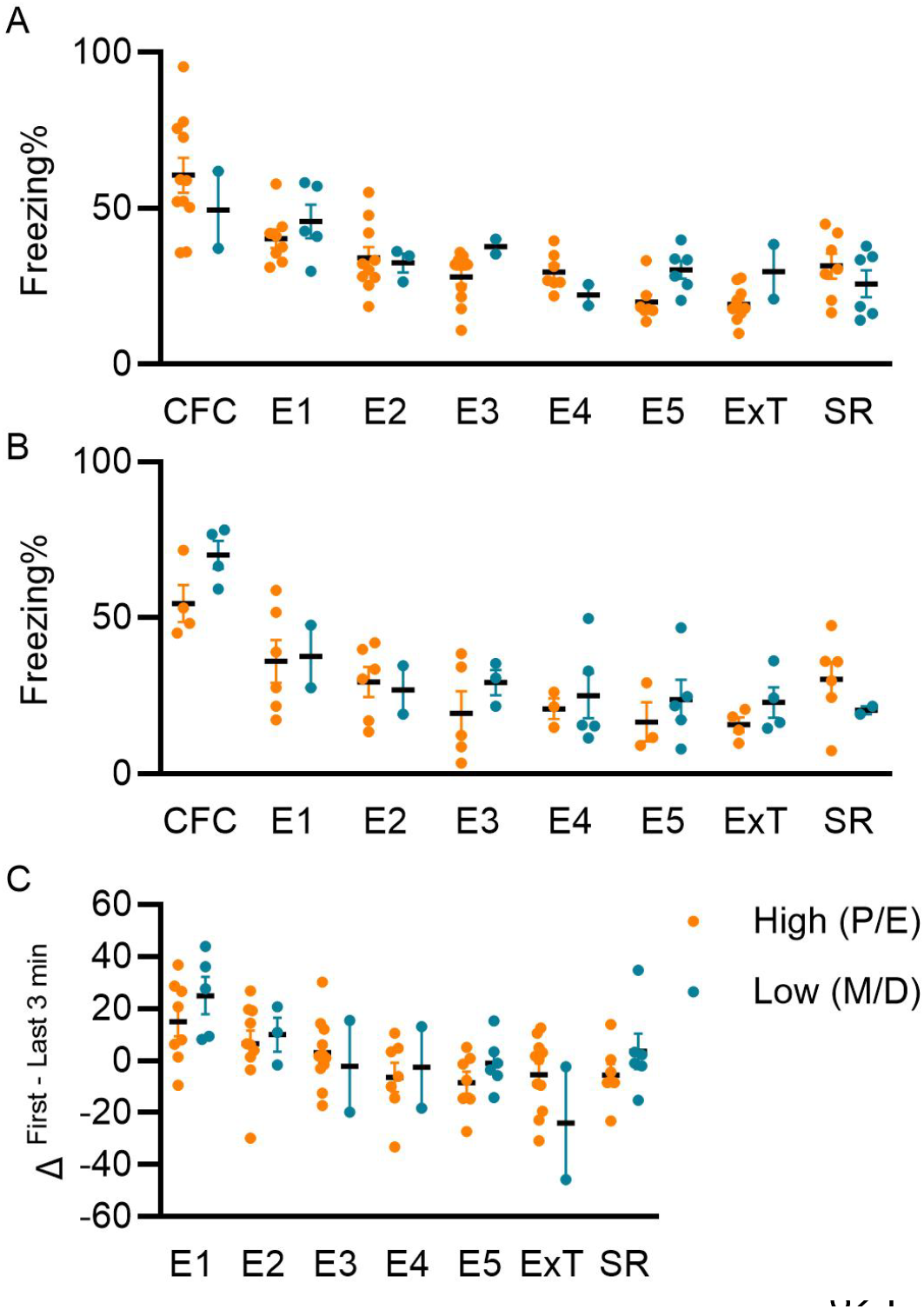
Estrous-stage comparisons of contextual fear conditioning, extinction training, and memory testing. Females were grouped according to estrous stage at each behavioral assessment into High (proestrus/estrus, P/E) and Low (metestrus/diestrus, M/D), reflecting stages broadly associated with higher and lower ovarian hormone levels, respectively, as described previously (Bouchet et al., 2017; Hohorst et al., 2025). **A)** Percentage of freezing during contextual fear conditioning (CFC), five consecutive extinction training sessions (E1-E5), the extinction memory test (ExT), and the spontaneous recovery test (SR) in females subjected to 6-minute daily extinction training (n = 13). **B)** Percentage of freezing during CFC, E1–E5, ExT, and SR in females subjected to 3-minute daily extinction training (n = 8). **C)** Within-session freezing reductions during 6-minute extinction training (E1-E5), calculated as the percentage freezing during the first 3 minutes minus that during the last 3 minutes of each session. No significant differences were detected between P/E and M/D groups across the behavioral assessments (Welch’s unpaired *t*-tests, corrected for multiple comparisons across sessions within each panel using the two-stage step-up false discovery rate method of Benjamini, Krieger and Yekutieli, Q = 1%; all *q* ≥ 0.15). In the 6-minute cohort, uncorrected *p*- values were 0.048 at E3 and 0.019 at E5 (*q* = 0.19 and 0.15, respectively), but neither was significant after correction. Data are presented as individual values with mean ± SEM. Sample sizes varied across assessments and estrous-stage groups (A, C: P/E n = 7-11, M/D n = 2-6; B: P/E n = 3-6, M/D n = 2-5).

